# Validation of an RNase H2 activity assay suitable for clinical screening

**DOI:** 10.1101/2022.06.09.494809

**Authors:** Marian Schulz, Claudia Günther, Rayk Behrendt, Axel Roers

## Abstract

As the key enzyme mediating ribonucleotide excision repair, RNase H2 is essential for the removal of single ribonucleotides from DNA in order to prevent genome damage. Loss of RNase H2 activity directly contributes to the pathogenesis of autoinflammatory and autoimmune diseases and might further play a role in ageing and neurodegeneration. Moreover, RNase H2 activity is a potential diagnostic and prognostic marker in several types of cancer. Until today, no method for quantification of RNase H2 activity has been validated for the clinical setting. Herein, validation and benchmarks of a FRET-based whole-cell lysate RNase H2 activity assay are presented, including standard conditions and procedures to calculate standardized RNase H2 activity. Spanning a wide working range, the assay is applicable to various human cell or tissue samples with overall methodological assay variability from 8.6% to 16%. The assay readily detected reduced RNase H2 activity in lymphocytes of a patient with systemic sclerosis carrying a *RNASEH2C* variant. Implementation of larger control groups will help to assess the diagnostic and prognostic value of clinical screening for RNase H2 activity in the future.

## Introduction

An increasing number of inflammatory and degenerative diseases are found to be associated with compromised genome integrity. In some of these, genome damage is assumed to be a central pathogenic event, while in others DNA damage may represent an epiphenomenon ^1–5^. Resolution of RNA/DNA hybrids is central to various DNA transactions and maintenance of genome integrity. Mammals express RNases H1 and H2 which both cleave RNA/DNA hybrids by catalysing phosphodiester bond hydrolysis ^6^. The enzymes play a role in resolution of R-loops, maturation of Okazaki fragments, repression of endogenous retroelements and degradation of RNA/DNA hybrids during cell death ^7–9^. While RNase H1 requires hybrids with at least 2 consecutive ribonucleotides, RNase H2 also cleaves single ribonucleotides embedded in the DNA double helix ^10^. Ribonucleotides are incorporated into genomic DNA in very high numbers during replication due to the limited capacity of the replicative polymerases to discriminate them from desoxyribonucleotides (Lujan et al., 2013; Sassa et al., 2019). To prevent destabilization of DNA, ribonucleotides are rapidly removed post-replication by the ribonucleotide excision repair (RER) pathway initiated by RNase H2-mediated nicking 5’ of the ribonucleotide. RER is the only error-free pathway capable of removing single ribonucleotides from DNA ^7–9^. Failure to repair these lesions leads to DNA damage ^8,11–16^. In mammals, complete loss of RNase H2 activity leads to embryonic lethality ^12,14^. Partial loss of function, however, caused by hypomorphic *RNASEH2* alleles can lead to autoinflammation and autoimmunity, as for example in Aicardi-Goutières syndrome (AGS), a monogenic ‘type I interferonopathy’ ^17–19^. Hypomorphic *RNASEH2* alleles also contribute to the polygenic predisposition for systemic lupus erythematosis (SLE) ^20–22^. Investigation of RNase H2-deficient human cells and mice recently led to elucidation of an important link between genome damage and chronic inflammation ^12^. DNA lesions ensuing from unrepaired ribonucleotides result in chromosomal aberrations, problems of mitotic segregation of defective chromosomes and formation of micronuclei. Upon collapse of the unstable micronuclear envelope, micronuclear chromatin is sensed by the intracellular DNA sensor cGAS, in turn resulting in activation of the sensor STING and activation of type I IFN and proinflammatory cytokine responses ^23–25^. Chronic activation of cGAS/STING signalling leads to autoinflammation, loss of T cell and B cell tolerance and autoimmune pathology ^18^. RNase H2 deficiency also predisposes to cancer in mice ^26,27^ and RNase H2 loss-of-function mutations occur in large fractions of human chronic lymphocytic leukaemia and prostate cancer ^28^. Reduced expression of RNase H2 is associated with reduced survival in colonic cancer ^27^. Conversely, upregulation of RNase H2 subunits was found to be a malignancy factor in numerous carcinomas and sarcomas ^29–31^. Moreover, double-strand breaks as resulting from compromised RNase H2 function were reported to contribute to neurodegeneration and aging ^32–34^. Collectively, RNase H2 is a relevant diagnostic and prognostic factor in diverse human disease settings, warranting clinical testing for RNase H2 activity in human cells or tissues.

Human RNase H2, unlike its monomeric prokaryotic isoenzyme RNase HII, is a heterotrimeric complex consisting of three proteins, the catalytic subunit RNase H2A and two auxiliary subunits, RNase H2B and RNase H2C ^35–37^. About 50 disease-causing *RNASEH2* variants have been identified to date ^21,22,24,38^, most of which are located in subunit B. While many variants exibit reduced RNase H2 substrate binding and hydrolysis, other mutant proteins did not show impaired activity in cell free assays using recombinant enzyme ^39^. The latter might feature compromized complex stability or interaction with additional proteins *in vivo*.

Although measurement of RNase H activity in mammalian cell samples has been performed since it’s discovery in 1969 ^40^, a standardized and validated method available for clinical use has been lacking. RNase H activity can be quantified by several different approaches relying on acid-insoluble precipitation, gel electrophoresis or HPLC. Two groups developed a fluorescence assay suitable for high-throughput studies and superior to earlier approaches with respect to precision, speed, labour and cost. RNase H2-mediated cleavage of a double-stranded DNA substrate containing a single ribonucleotide results in release of a fluorescein-labelled fragment from a quencher ^41,42^. Herein, we adapt this assay into a standardized and validated procedure relying on whole cell lysates for clinical screening of effective intracellular RNase H2 activity.

## Materials and Methods

### Ethics approval and control group selection

Ethics approval was granted by the ethics committee of the Medical Faculty Carl Gustav Carus, TU Dresden (EK 31022012). Volunteers older than 18 years of age without overt disease for the past two weeks were included after informed consent. Pregnancy or medication, abuse of alcohol or drugs were exclusion criteria. Volunteers did not receive financial or other compensation.

### Cell culture

HeLa cells and murine embryonic fibroblasts (MEFs) were cultured in Gibco® DMEM – Dulbecco’s Modified Eagle Medium (Fisher Scientific GmbH, Schwerte, Germany) at 37 °C and 5% CO_2_. For harvesting, medium was aspirated, adherent cells were washed twice with 1x PBS followed by incubation with 1x trypsin (0.25%, Life Technologies Germany, Darmstadt, Germany) at 37°C for 2 minutes. Digest was stopped by addition of FCS-containing medium, cells were detached by pipetting, transferred into a 15-ml conical tube and pelletted at 330 x g for 5 min. Cells were resuspended, washed twice in 5 ml of chilled PBS for freezing, or in 1x FACS buffer for FACS sorting. For freezing, supernatant was discarded, pellets were shock-frozen in liquid nitrogen and then stored at -80°C for a maximum of 4 weeks.

### Isolation of Primary cells from human blood and mice

For isolation of human PBMC, peripheral blood was collected in 10 ml heparinized tubes, stored at 4°C and analysed within 4 hours. Blood was diluted in an equal volume of PBS (calcium- and magnesium-free, equilibrated to room temperature (RT)). PBMC were isolated by standard Ficoll®-Paque density gradient centrifugation, and washed 3 times with chilled PBS. Murine keratinocytes, peritoneal cells, splenocytes and embryonic fibroblasts were isolated by standard procedures ^43–46^

### Flow cytometric cell sorting

PBMC from human donors were stained with anti-human CD3 (UCHT1) PE and anti-human CD19 (SJ25-C1) APC-H7, murine spleen cells with anti-CD19 (eBio1D3) PE, anti-CD4 (RM4-5) APC, anti-CD11b (M1/70) eF450 and anti-CD11c (N418) PE/Cy7, murine peritoneal lavage cells with anti-CD11b (M1/70) eF450 and anti-F4/80 (BM8) PE, and murine epidermal cells with anti-CD49f (eBioGoH3 rat) PE antibodies at 4°C for 30 minutes. Antibodies were purchased from Thermo Fisher Scientific Germany (Frankfurt a. M., Germany). Stained cells, or harvested cell culture cells, respectively, were washed with FACS buffer three times and resuspended in FACS buffer. Shortly before analysis, DAPI was added to a final concentration of 3 μM. Cells were sorted on a BD FACSAria^™^ III (Beckton Dickinson Germany, Heidelberg, Germany) excluding doublets and dead cells. Data was analyzed using FlowJo Single Cell Analysis Software (FLOWJO, LLC Data analysis software).

### Cell lysis and protein quantification

Washed cell pellets were dissolved in a suitable amount of lysis buffer 1 and incubated on ice for 10 min. After addition of the same amount of lysis buffer 2 and another incubation on ice for 10 minutes, cell debris was spun down at 20 000 x g for 10 min at 4°C. Supernatant containing total cellular protein was harvested, and replicates were stored at -80°C. Protein concentration was determined using the Qubit^™^ Protein Assay Kit (ThermoFisher scientific) following recommendations of the vendor.

### RNase H2 activity assay and standard conditions

RNase H2 activity was measured using a fluorometric assay approach adapted from Crow et al. (Crow et al., 2006). The type 2 RNase H-specific substrate consisted of an 18 bp DNA strand containing a single ribonucleotide 4 bp 5’ of a covalently attached 3’ fluorescein residue (oligonucleotide B) which was annealed to a 18 bp anti-sense DNA strand with a quenching 5’ dabcyl residue (oligonucleotide D). Type 2 RNase H hydrolyses the phosphodiester bond 5’ of the single ribonucleotide leading to dissociation of the fluorescein-carrying fragment from the quencher allowing photometric quantification (Figure 1A). As positive controls, unquenched single-stranded substrate (B), unquenched double-stranded substrate lacking the dabcyl residue at the anti-sense strand (BK), and plateau-fluorescence of the fluorescence progress curve (BD plateau), were implemented. BD plateau-fluorescence was determined by measuring fluorescence in wells containing 100 eqU HeLa RNase H2 and different amounts of substrate BD for 270 min until there was no further fluorescence increase. The mean of the last three measurement value triplicates was defined as BD plateau-fluorescence. As negative controls, quenched type 2 RNase H-specific substrate without addition of RNase HII (BD), quenched type 2 RNase H-specific substrate with addition of *heat-inactivated* cell lysate (BD + *h*.*i. lysate*), quenched double-stranded 2-O’-methylated RNA / DNA (type 2 RNase H-resistant) substrate (AD) with addition of active RNase HII, and blanks, were used. Desalted oligonucleotides were purchased from Eurogentec (Seraing, Belgium), dissolved in TE buffer to a final concentration of 100 μM and annealed by heating to 90 °C for 2 minutes and then gradually cooling down by 1 °C per minute. Then, substrates were aliquoted and stored at -20 °C at a concentration of 10 pmol / μl.

**Figure 1.**
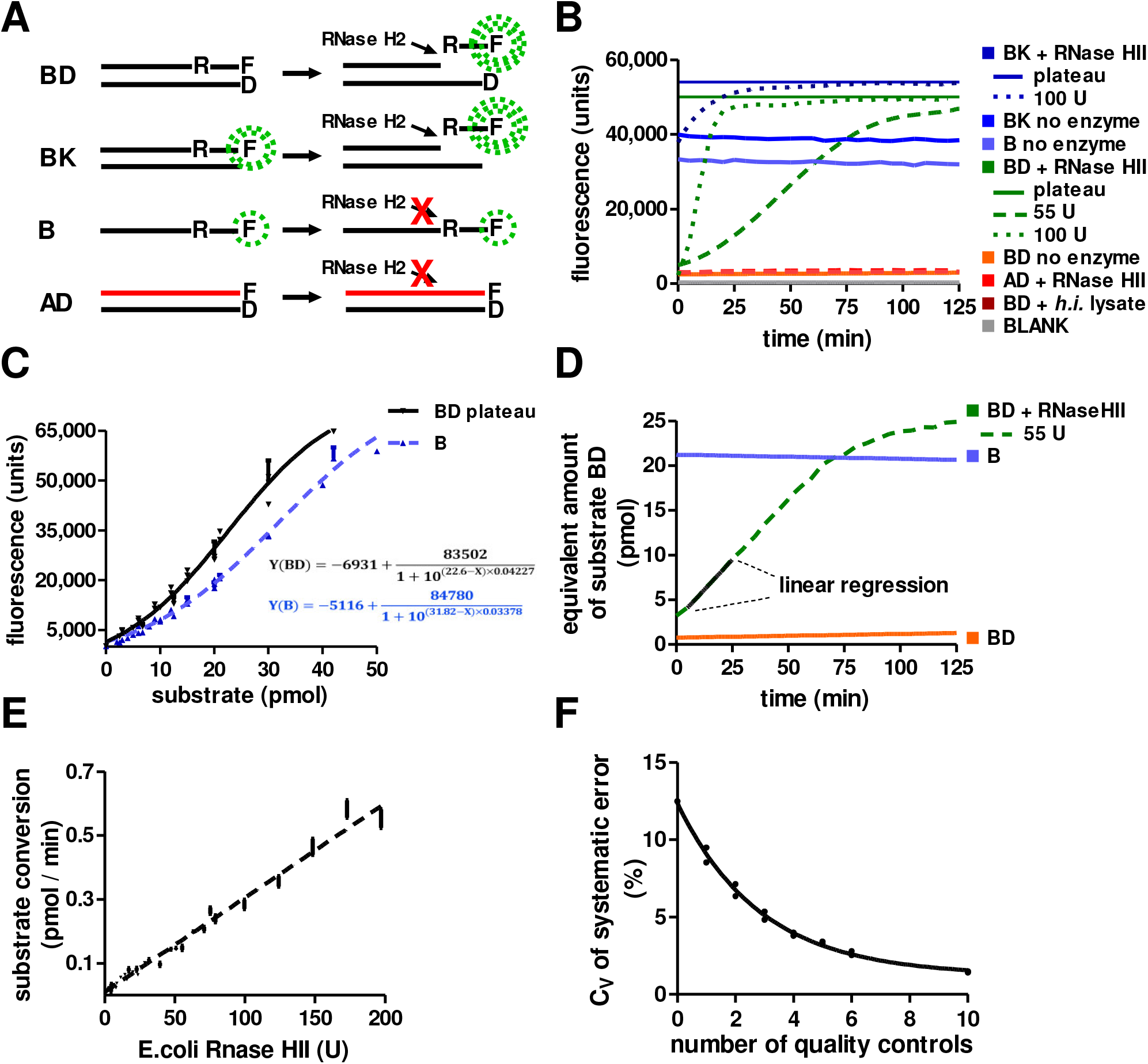
Assay principle and implementation of controls. **A)** RNase H2 activity assay design (adapted from Crow et al. ^42^). Substrate BD, specific for mammalian RNase H2 and bacterial RNase HII, is an 18 bp double-stranded DNA containing a single ribonucleotide and a fluorescent residue (fluorescein) as well as a quencher (dabcyl) on the complementary strand. RNase H2 cleavage at the position of the ribonucleotide releases fluorescein from the quencher. Positive control substrates BK and B are identical but lack the quencher or the complementary strand, respectively. RNase H2 does not cleave single-stranded substrates like substrate B. Negative control substrate AD contains methylated RNA resistant to RNase H2 and was used to control for substrate dequenching by spontaneous dissociation. **B)** Addition of 100U RNase HII lead to cleavage of positive control BK as well as substrate BD with fluorescence reaching a plateau (BK plateau and BD plateau) above the fluorescence levels of positive controls B and BK. This aligns with a quenching effect of the complementary strand ^56^. Positive control BK exceeded positive control B fluorescence when no enzyme was added. This is explained with decreased fluorophore stability of substrate B lacking the protective effect of the quenching complementary strand during storage ^57^. Thus, BD plateau fluorescence is the only valid positive control for calculation of substrate cleavage. Fluorescence negative controls included quenched double-stranded substrate BD without addition of RNase HII (BD no enzyme) and with addition of *heat-inactivated* cell lysate (BD + *h*.*i*. lysate), quenched type 2 RNase H-resistant substrate (AD + RNase HII) and blanks. Negative control fluorescence reached a maximum of 6.06 % (95 % CI: 5.26 % - 6.86 %) of BD plateau fluorescence. Unspecific substrate cleavage or degradation was insignificant with 0.09 % per minute (95 % CI: 0.03 % / min - 0.15 % / min). Fluorophores showed stable fluorescence with a fluorescence decrease of 1.68 % per hour (95%CI: 0.59 % / h - 2.78 % / h). Addition of lower amounts of RNase HII (e.g. 55 U, other curves are not shown) yielded an increase in fluorescence with unexpectedly long lag phase before a hardly definable steady state phase and a plateau phase. **C)** Implementation of fluorescence standard curves revealed concentration-dependent fluorescence non-linearity of the fluorophores. B and BD plateau fluorescence were measured in triplicates at eight different substrate concentrations (20nM - 500nM) after addition of 100 U RNase HII. **D)** With help of the BD plateau fluorescence standard curve, fluorescence progress curves were transformed into substrate cleavage progress curves now showing a perfectly linear segment indicating steady-state conditions of the pseudo-first order irreversible cleavage reaction. **E)** A substrate conversion standard curve was implemented using different amounts of E.coli RNase HII. Using this curve and quality controls, measured catalytic activity can be assigned a standardized, externally validated unit (“eqU”). The curve showed no significant deviation from linearity in the implemented assay working range (8.09 U – 200 U) (linear regression r square = 0.99; Run’s test: deviation from linearity not significant, P = 0.44). **F)** Mean systematic error between separate RNase H2 assays performed under the same assay conditions was 3.65 % (SD = 1.78; n = 3). The precision by which this systematic error could be calculated was dependent on the number of quality controls used. Use of six matched quality controls reduced variability of the calculation of systematic error to less than 3%. Mean plus/ minus SD is shown.

After cell isolation, cell number or protein content was quantified and cell pellets were lysed as described above. Cell lysates were premixed on ice with an 1 : 1 mixture of lysis buffer 1 and 2 in a 96-well flat-bottomed plate. Then, equal amounts of cell lysate premixes were pipetted to another flat-bottomed 96-well reaction plate containing 100 μl reaction buffer with 270 nM type 2 RNase H-specific substrate (BD) using a multi-channel pipette. The reaction was monitored in a FLUOstar® Omega photometer at 37 °C for at least 120 min, fluorescence was measured at 3 minute intervals. Before measurement, the photometer was calibrated, setting the 30 nM unquenched single-stranded substrate B positive control fluorescence to 33333 FU. Photometer measurement range was set to 100 %. A 485 nm excitation filter and a 520 nm emission filter were used. Before each measurement, wells were mixed by orbital shaking (3 mm diameter, 5 s). Fluorescence was induced by 10 flashes per well and cycle, and measurement was performed by orbital scanning by a vertically adjusted sensor. Fluorescence was measured using a time-resolved approach with an integration delay of 47 μs and an integration time of 1510 μs. Positioning delay was set to 0.2 s, measurement start time to 0.5 s. All assay steps except from photometric measurement were carried out on ice.

Fluorescence data was converted into the equivalent amount of cleaved substrate BD using the BD plateau fluorescence standard curve to acquire substrate cleavage progress curves. The cleavage rate was obtained by linear regression of the curves between minute 3 and 24 (at least 5 data points). Using six quality controls with known activity and the substrate conversion standard curve (Figure 2D), cleavage rates were transformed into standardized catalytic activities (1 “eqU” = equivalent to the catalytic activity of 1 U RNase HII (NEB ®) under standard conditions). To correct for systematic error between different RNase H2 assays, six internal standards (cell lysate aliquots with known RNase H2 activity) were measured in each experiment. For oligonucleotide sequences and buffer reagents, see Table 1 and Table 2.

**Table 1.**

| buffer/ solution | reagents |
| --- | --- |
| RNase H2 assay reaction buffer (1x) | 60 mM KCl, 50 mM Tris.HCl pH 8.0, 20 mM MgCl<sub>2</sub>, add fresh Triton X-100 and BSA to a final concentration of 0.01% |
| lysis buffer 1 (1x) | 50 mM TRIS.HCL pH 8.0, 280 mM NaCl, 0,5% v/v NP40, 0,2 mM EDTA, 0,2 mM EGTA, 10% v/v glycerol, 0.1 mM sodium orthovanadate, add fresh 1 mM DTT and 1mM PMSF |
| lysis buffer 2 (1x) | 20 mM HEPES, 10 mM KCl, 1 mM EDTA, 0.1 mM sodium orthovanadate, add fresh 1 mM DTT and 1mM PMSF |

**Table 2.**
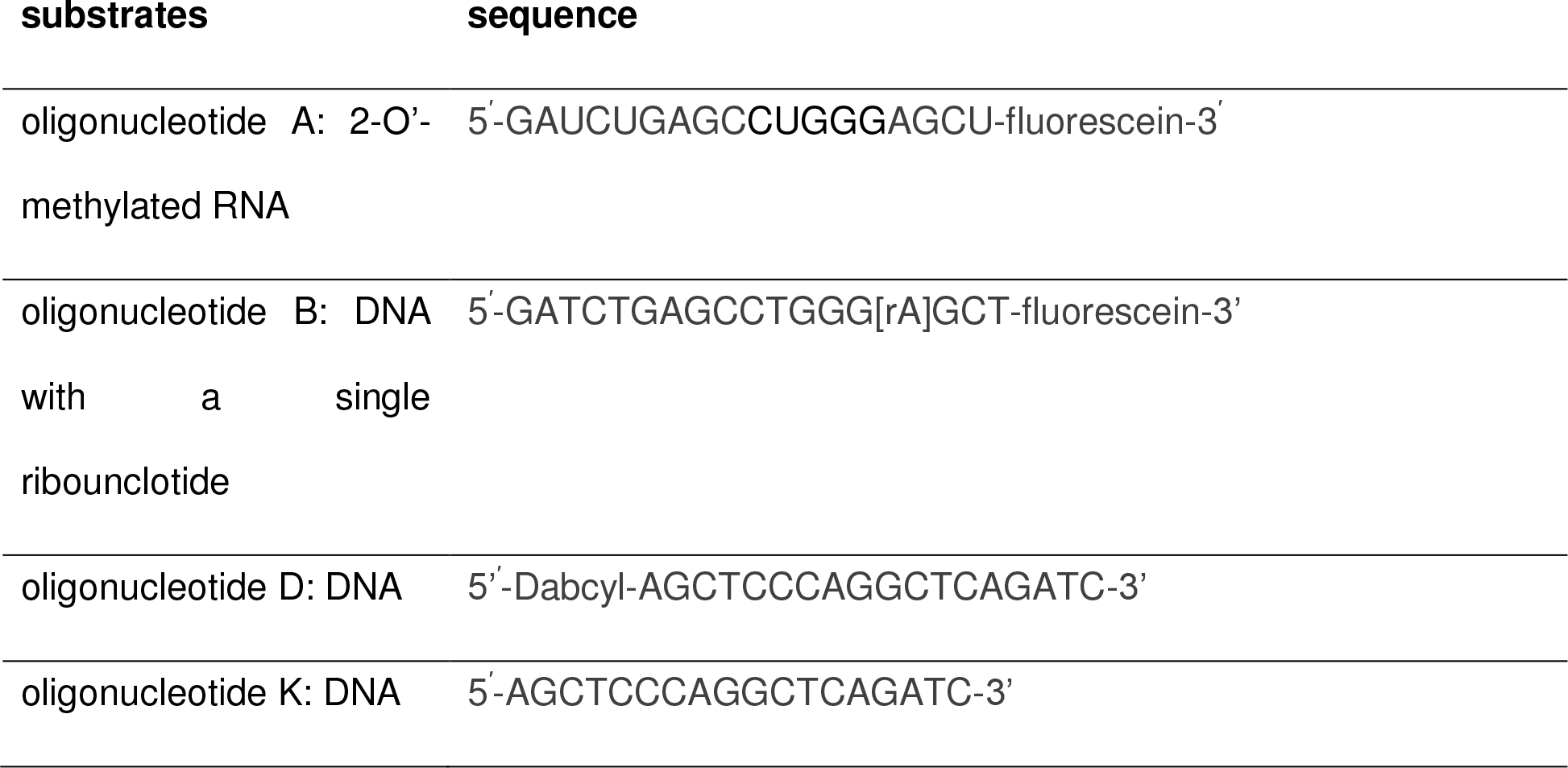
RNase H2 assay.

**Figure 2.**
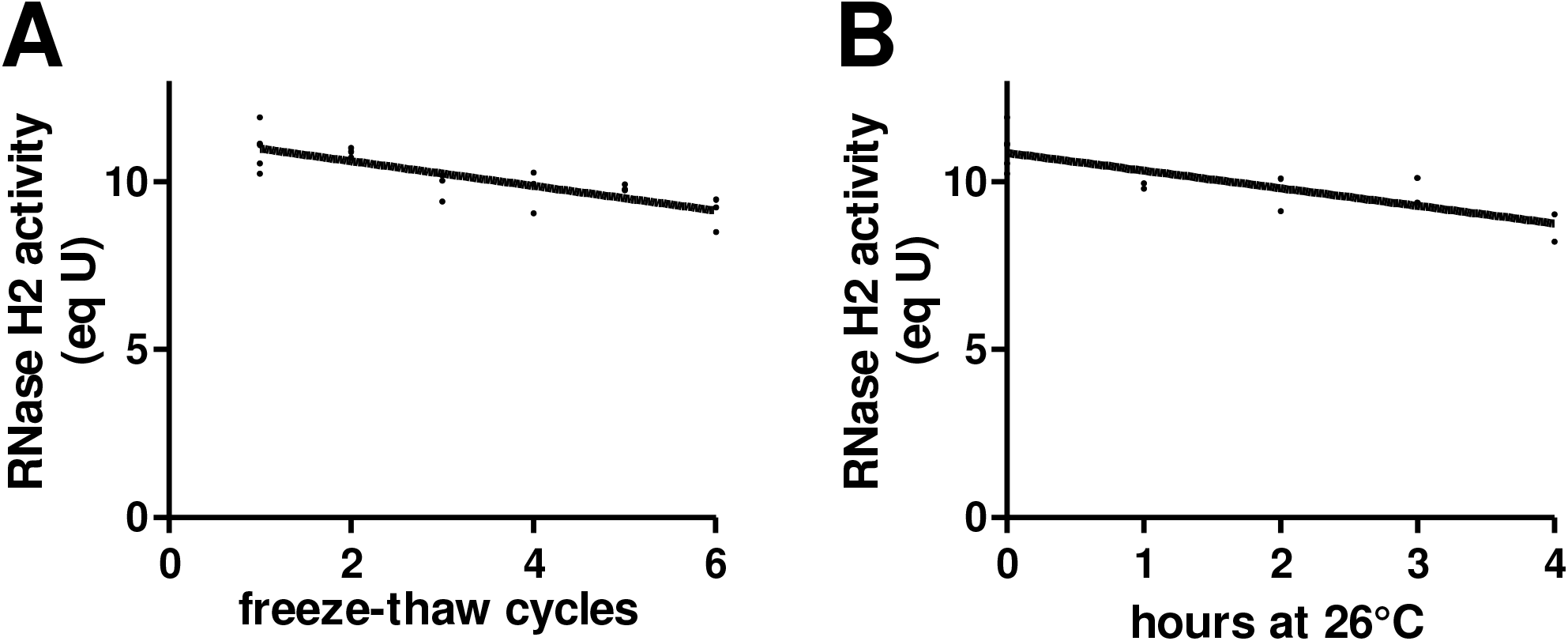
Ruggedness. **A)** Aliquots of the same sample were frozen at -20 °C and thawed up to 6 times. Each cycle of freezing and thawing resulted in a mean activity loss of -0.37 eqU (Pearson’s r = -0.94). **B)** Aliquots of the same sample were stored at 26°C for up to 4 hours and activities were compared with the RNase H2 assay. Mean loss of activity was -0.53 eqU/ hour (Pearson’s r = -0.91).

### Statistics

Statistical tests were performed using GraphPad Prism^™^ 5.04 (GraphPad Software Inc., San Diego, California, USA). Sample size calculations (two-sample t-test) were conducted as proposed by Hulley et al. ^47^ using G*Power Version 3.1.9.4 © (Franz Faul, Kiel, Germany).

## Results

### Validation and benchmarks of the RNase H2 activity assay

The assay principle was adapted from Crow et al. ^42^ as illustrated in Figure 1A. A double-stranded DNA oligonucleotide containing a single ribonucleotide and a fluorescent label (fluorescein) at the 3’ end of the same strand were used as a type 2 RNase H-specific substrate. This substrate is cleaved by mammalian RNase H2, but also by bacterial RNase HII. Fluorescence was quenched by a dabcyl-group coupled to the 5’ end of the complementary strand. Upon RNase H2-mediated cleavage at the position of the ribonucleotide, a short oligonucleotide carrying the fluorescein label is released from the quencher and fluorescence is quantified by photometry.

### Implementation of controls and standard curves

Figure 1B shows fluorescence progress curves for type 2 RNase H-specific (BD) and control substrates (B, BK, AD, see Figure 1A). Bacterial RNase HII cleaved substrate BD with fluorescence reaching a plateau (Figure 1B), while this substrate showed only weak spontaneous background fluorescence (6.06% of plateau level). Spontaneous dequenching by degradation of quenched substrate BD was insignificant (Figure 1B). Addition of heat-inactivated cell lysate had no effect on fluorescence, indicating absence of unspecific (heat-sensitive) quenchers. Likewise, there was no unspecific substrate degradation detectable upon addition of active cell lysate to type 2 RNase H-resistant substrate AD (Figure 1B). Maximum fluorescence of fully cleaved substrate BD (BD plateau fluorescence) was defined as high control for substrate conversion.

To allow inter-laboratory reproducibility, the assay was validated using E. coli RNase HII with standardized activity. Different amounts of RNase HII were added to samples containing substrate BD and fluorescence progress curves (results for 55 U and 100 U RNase HII shown in Fig. 1B) were determined in pipetting triplicates under standard assay conditions. Fluorescence progress curves were double-curved exhibiting a significant lag phase. This was unexpected for a pseudo-first order irreversible reaction with a single substrate and without any known inhibitors or any described conformational changes of the enzyme during the reaction ^48,49^. Therefore, it was hypothesized, that this lag phase was due to non-linear, concentration-dependent fluorescence behaviour of the fluorophore. Indeed, implementation of a positive control fluorescence standard curve by enzymatic dequenching of substrates BD and B demonstrated fluorescence non-linearity of the fluorophore (Figure 1C). We therefore converted fluorescence data into amount of cleaved substrate based on the fluorescence standard curve (Figure 1C). Increase of amount of cleaved substrate over time showed perfectly linear behaviour without a significant lag phase (Figure 1D) allowing for definition of the steady-state phase, in which linear regression was performed to calculate RNase H2 activity. Linearity was highly significant with an r square value of >0.99 for all curves corresponding to RNase HII activity above 8.09 U (limit of quantification (LOQ), see below). Hence, steady state conditions could be assumed for enzyme activities between 8.09 and 200 U of E. coli RNase HII under standard conditions.

Substrate B positive control fluorescence was used for photometer calibration to ensure reproducibility of experimental data. Plotting catalytic activities of standardized amounts of RNase HII yielded a substrate conversion standard curve (Figure 1E). Using this standard curve, measured catalytic activity was converted into the equivalent activity (“eqU”) of a defined amount of externally validated reference-RNase HII. Substrate conversion rates showed no significant deviation from linearity in the validated working range indicating small systematic error as well as absence of activators or inhibitors in the reaction mix.

For comparison and combination of data from different RNase H2 assays, internal standards (see Materials and Methods) were used. *Inter*-assay systematic error was assessed between three individual RNase H2 activity assays using aliquots of the same mouse embryonic fibroblast lysate. Means differed by 3.65% (SD=1.78; n=3). However, the precision by which this systematic error was calculated was strongly dependent on the number of matched quality controls established between the individual assays (Figure 1F). Use of six matched quality controls reduced variability of the calculation of systematic error to less than 3%.

### Sensitivity and ruggedness

The average curve slope of substrate BD without addition of enzyme was 7.8 fluorescence units (FU) per minute (0.026 % of positive control fluorescence, calculated from data shown in Figure 1B). This was equivalent to a substrate cleavage rate of 5.4 fmol/min resulting in a limit of detection (LOD) of 9.7 fmol/min (2.09 eqU) and an LOQ of 32.3 fmol/min (8.09 eqU)^50^.

Designing the RNase H2 assay for high-throughput analysis required prolonged sample handling and sample storage. Systematic sample handling error was assessed by subjecting HeLa whole cell lysate to repeated freezing and thawing (Figure 2A) or by incubation at RT for defined time (Figure 2B). Each freeze-thaw cycle reduced RNase H2 activity by 0.37eqU (4.5% of LOQ). Likewise, incubation at RT reduced enzyme activity at a rate of 0.53 eqU per hour (6.6% of LOQ).

Standard sample handling involved one freeze-thaw cycle and incubation times between 30 minutes and 2 hours, which were, however, performed on ice rather than RT, suggesting maximal loss of RNaseH2 activity of 1.43 eqU (17.7 % of LOQ) due to the processing.

### Steady-state kinetics and assay endpoints

In search for the most-suited parameter to be determined (assay endpoint) for purposes of clinical screening, RNase H2 steady-state kinetics was studied. For this, 2.5 μg of HeLa protein (30 eqU) were added to wells containing eight different substrate concentrations spread around the expected K_M_-value ^6,51^. This was performed with HeLa protein from six individual HeLa cell cultures. Michaelis-Menten curves were determined using the initial-rates method and Michaelis-Menten non-linear regression (Figure 3). RNase H2 activity followed Michaelis-Menten kinetics with a mean K_M_ of 141.7 nM and a high mean C_V_ of 65.72 %. Calculation of V_MAX_ was less variable with a mean C_V_ of 24.45%. Still, highest precision was obtained by measuring RNase H2 activity at a single substrate concentration (provided a substrate concentration >2SD above K_M_ was used). At a substrate concentration of 270 nM, RNase H2 activity reached approximately 60% of V_MAX_ and measurement variability was below 10%. Higher substrate concentrations lead to a systematic distortion of RNase H2 activity due to non-linear fluorescence behaviour of the fluorophores (Figure 1C). Therefore, it was concluded, that RNase H2 activity at a substrate concentration of 270 nM was best-suited as assay endpoint for purposes of clinical screening.

**Figure 3.**
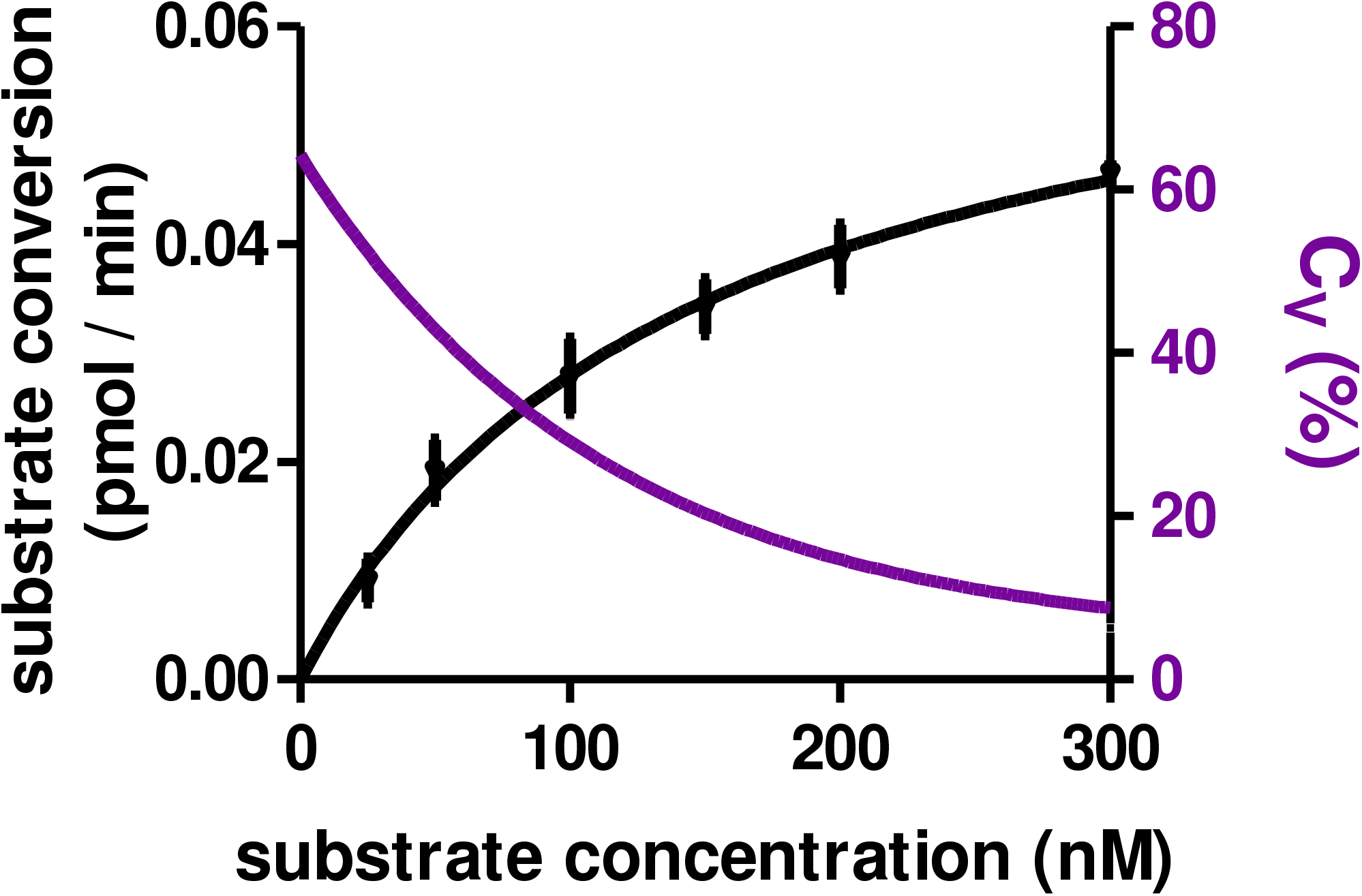
Steady-state kinetics. The mean K_M_ was 141.7 nM (SEM = 27.86 %; 95 % CI: 69.4 – 213.9 nM). Calculation of K_M_ and V_MAX_ values showed high variability (C_V_ (K_M_) = 65.72 %; C_V_ (V_MAX_) = 24.45 %), while the C_V_ of individual measurements was dependent on the substrate concentration. RNase H2 activity at a substrate concentration of 270 nM (> 2 SD above K_M_) was associated with a C_V_ below 10 % and implemented as assay end-point. Mean ± SD is shown.

### Precision

Overall precision was determined for the assay based on primary or cultured cells. Hereby, the influence of different assay steps (cell isolation and preparation, pipetting, photometric measurement and linear regression) and normalization methods (normalization to cell number or total protein) on overall variability was assessed. Total assay variability including all biological and methodological error sources ranged from 8.6% to 16% (C_V_).

Variability due to linear regression, photometer imprecision or pipetting was assessed in an experiment with 105 pipetting replicates and averaged at 7.73 %. This accounted for more than half of total assay variability (56.2 %; 95 % CI = 14.7 % - 97.8 %, n = 4), depending on the experimental approach and normalization method (contribution of single error sources on the total coefficient of variation was calculated using addition of variances). Under standard conditions, the largest part of this methodological error was attributed to pipetting error, while linear regression and photometer imprecision constituted minor error sources (not shown). Assay precision was strongly dependent on the normalization method used. Experiments relying on normalization to total protein avaraged a total assay variability of 9.6 %, unaffected by the cell isolation method (FACS-sorting or direct lysis of cultured cells), while normalization to cell number resulted in a much larger total assay variability of 16 % (Figure 4). We attributed this primarily to loss of cells during washing steps and, less predominantly, imprecisions of FACS cell counting. Isolation of primary cells from peripheral blood by Ficoll gradient centrifugation is known to yield PBMCs of variable cell type composition and viability ^52,53^. Thus, isolation of primary blood cells resulted in higher overall variability (11.17 %) than direct lysis of cultured cells.

**Figure 4.**
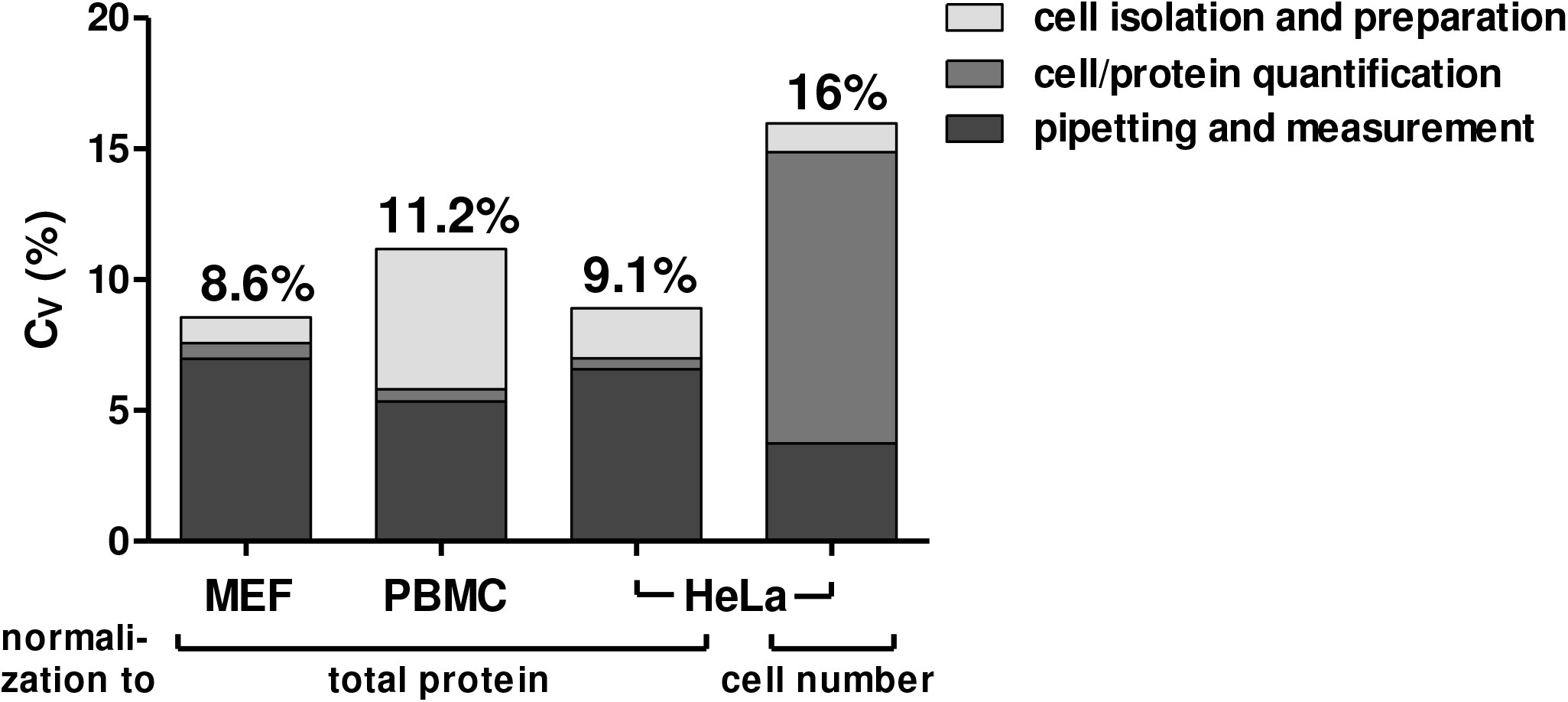
Assay precision. RNase H2 activity was determined in lysates of mouse embryonic fibroblasts (MEFs), lysates of PBMCs isolated from human blood by Ficoll-Paque gradient centrifugation, lysates of HeLa cells (sorted for living cells via FACS) with normalization to total protein or cell number as indicated. Total assay variability of the different approaches is shown above the bars. Stacked colours indicate contributions of different error sources (cell isolation and preparation; cell/protein quantification; pipetting and measurement imprecision determined in separate experiments (not shown).

### Screening RNase H2 activity in human lymphocytes

RNase H2 activity differed significantly between cell types (Figure 5). In mouse cells, RNase H2 activity changed with cell cycle phase (Supplemental Figure S2). To reduce variability, we aimed to assay for enzyme activity in one particular cell type rather than in samples containing undefinded mixtures of different cells. We chose to base the assay on lymphocytes as they are easily obtained in large numbers from blood and feature high RNase H2 activity (Figure 5 and Supplemental Table S1). PBMCs were obtained from blood samples of 24 healthy donors (Supplemental Figure S1) with unknown *RNASEH2* genotypes by FICOLL® gradient centrifugation. CD19^+^ B cells and CD3^+^ T cells were isolated by flow cytometric sorting. From a 10 ml blood sample, 4.0 × 10^5^ B cells and 3.0 × 10^6^ T cells were obtained, sufficient for multiple replicate measurements (Supplemental Table S1). Control group size was designed to allow detection of a reduction of RNase H2 activity by 30 % with a statistical power of 90 % and *α* of 0.10.

**Figure 5.**
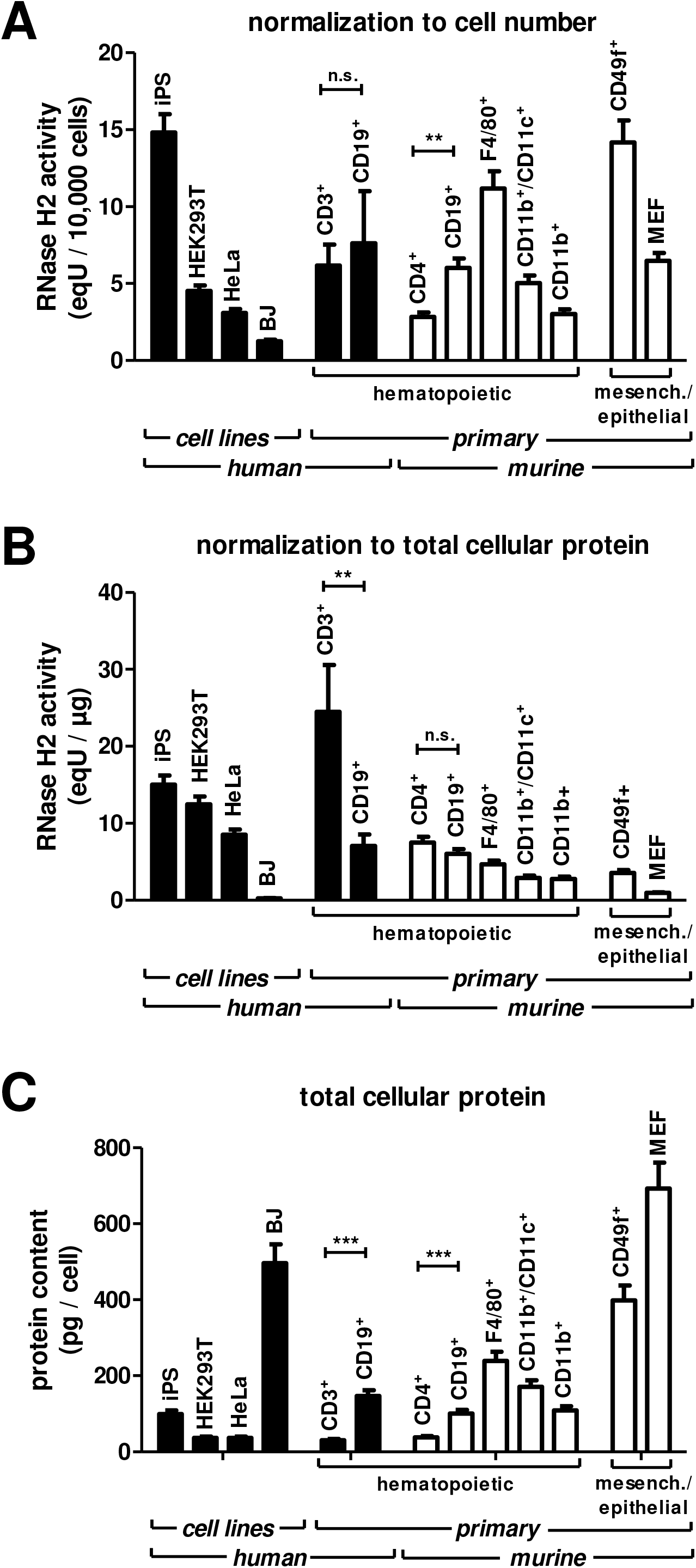
RNase H2 activity of different cell types. Human induced pluripotent stem (iPS) cells, human embryonic kidney 293 cells (HEK293T), HeLa cells, human fibroblasts from the BJ cell line, human peripheral blood T cells (CD3^+^) and B cells (CD19^+^), murine spleen T cells (CD4^+^), B cells (CD19^+^), dendritic cells (CD11b^+^/CD11c^+^) and macrophages (CD11b^+^), murine peritoneal macrophages (F4/80^+^), murine epidermal stem cells (CD49f^+^) and mouse embryonic fibroblasts (MEF) were purified and counted by flow cytometry. Cells were lysed, protein concentration was determined and RNase H2 activity was measured in biological triplicates under standard assay conditions. **A)** RNase H2 activity normalized to cell number. **B)** RNase H2 activity normalized to amount of total protein. **C)** total cellular protein content of the cell types. Mean ± SD is shown, significance was tested with the unpaired two tailed t-test, **** p < 0.0001, *** p < 0.001, ** p < 0.01 * p < 0.5, n.s. not significant.

In T cells, RNase H2 activity per μg of cellular protein was about 3-fold higher than in B cells (Figure 6B), while RNase H2 activity per cell did not differ significantly between B and T cells (Figure 6C), reflecting higher total protein content of B cells compared to T cells. *Inter-*individual assay variability in T cells was approximately four-fold lower as in B cells, irrespective of the normalization method (Fig. 6D).

**Figure 6.**
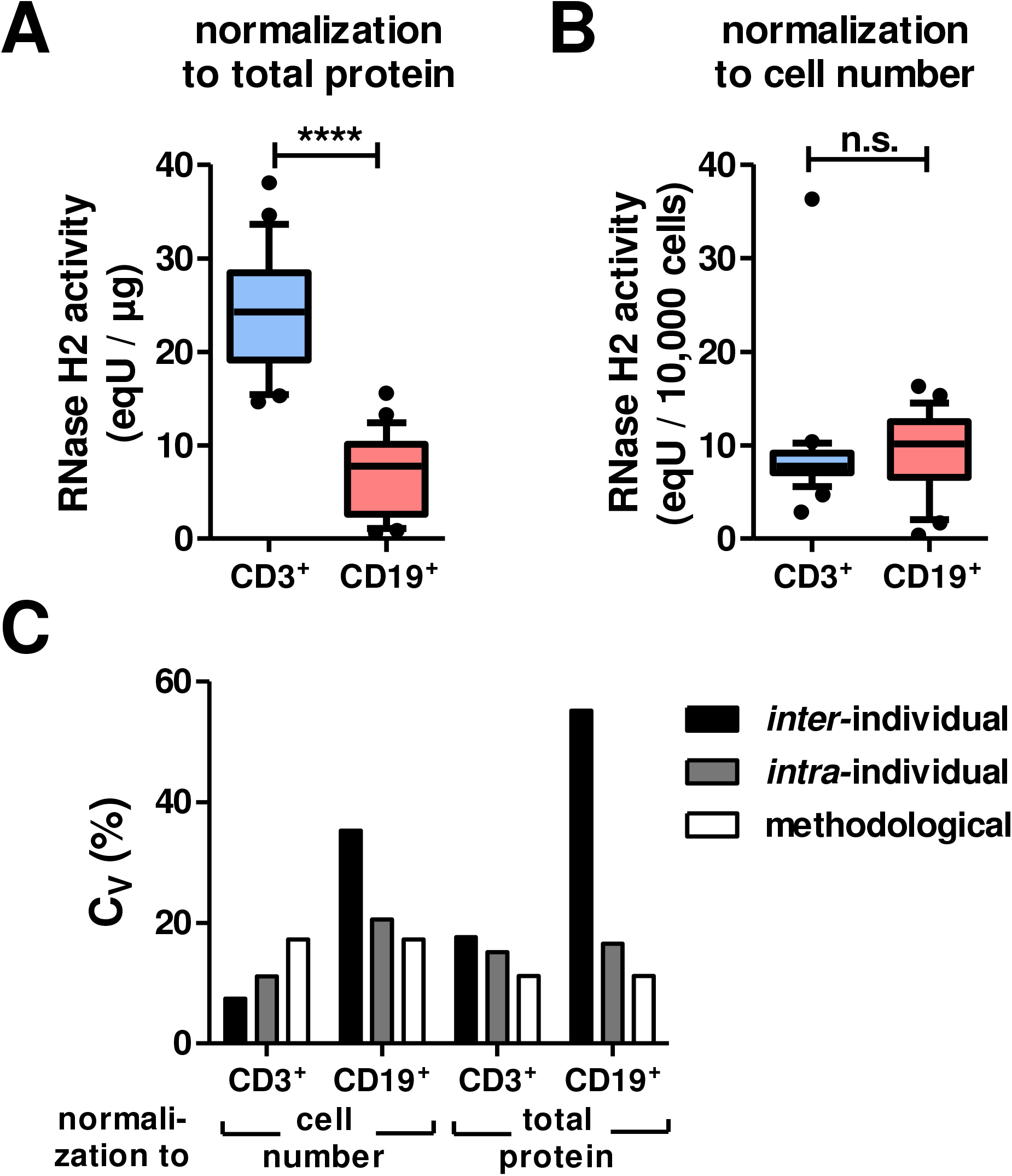
Control group benchmarks. **A)** RNase H2 activity normalized to total protein in T cells was significantly higher than in B cells. **B)** RNase H2 activity normalized to cell number did not differ significantly, but showed significantly smaller variability in T cells. **C)** Strong *inter-*individual variability was observed in B cells, while inter- and intra-individual variability were approximately on the same level as methodological error in T cells. Methodological variability was determined in validation experiments. Median is shown, boxes indicate 25^th^ and 75^th^ percentile, whiskers indicating 10^th^ and 90^th^ percentile. Extreme outliers (>95^th^ percentile oder < 5^th^ percentile) are shown in the diagram, but were excluded for statistical testing. Normality could be assumed for all groups (Shapiro-Wilk test). Significance was tested via unpaired two tailed t-test, **** p < 0.0001, *** p < 0.001, ** p < 0.01 * p < 0.5, n.s. not significant.

In T cells, *inter-* and *intra-*individual variability did not differ significantly. When activity was measured in T cells with normalization to cell numbers, the highest error source was methodological variability. In B cells, however, *inter*-individual variability clearly exceeded *intra-*individual and methodological variability. Gender and age did not contribute to *inter-*individual variability (not shown).

Collectively, we show that quantification of RNase H2 activity in T cells requires small amounts of venous blood and shows little *inter*-individual and *intra*-individual variation making it a suitable method for clinical use.

### *RNASEH2C c*.*468G>T* reduces RNase H2 activity in T cells

A pilot experiment was performed on a systemic sclerosis patient carrying an *RNASEH2C* variant with so far unknown effects on RNase H2 activity (c.468G>T, rs61736590). The condition of this 60-year old female was classified as ‘limited disease’ (onset with 33 years) with antinuclear antibodies (titre 1:2560, Scl 70), Raynaud’s syndrome, mutilation of finger tips by digital ulcers at onset of the disease, lung fibrosis and oesophageal involvement. She received no immunomodulatory or immunosuppressive therapy. The heterozygous mutation in *RNASEH2C* was identified by whole exome sequencing and verified by Sanger sequencing of DNA from PBMCs (Figure 7A). RNase H2 activity measured in triplicate samples was compared to the group of healthy controls (n = 24, Supplemental Figure S1) using unpaired two-tailed t-tests with Welch’s correction. RNase H2 activity per cell was significantly reduced in the patient’s T cells compared to the control group. Residual activity per cell and per μg of cellular protein in T cells was 80 % and 63.5 % of control activity, respectively (Figure 7B-D). Collectively, our data show that the assay readily detected the reduction of RNase H2 activity caused by the heterozygous *RNASEH2 c*.*468G>T* variant.

**Figure 7.**
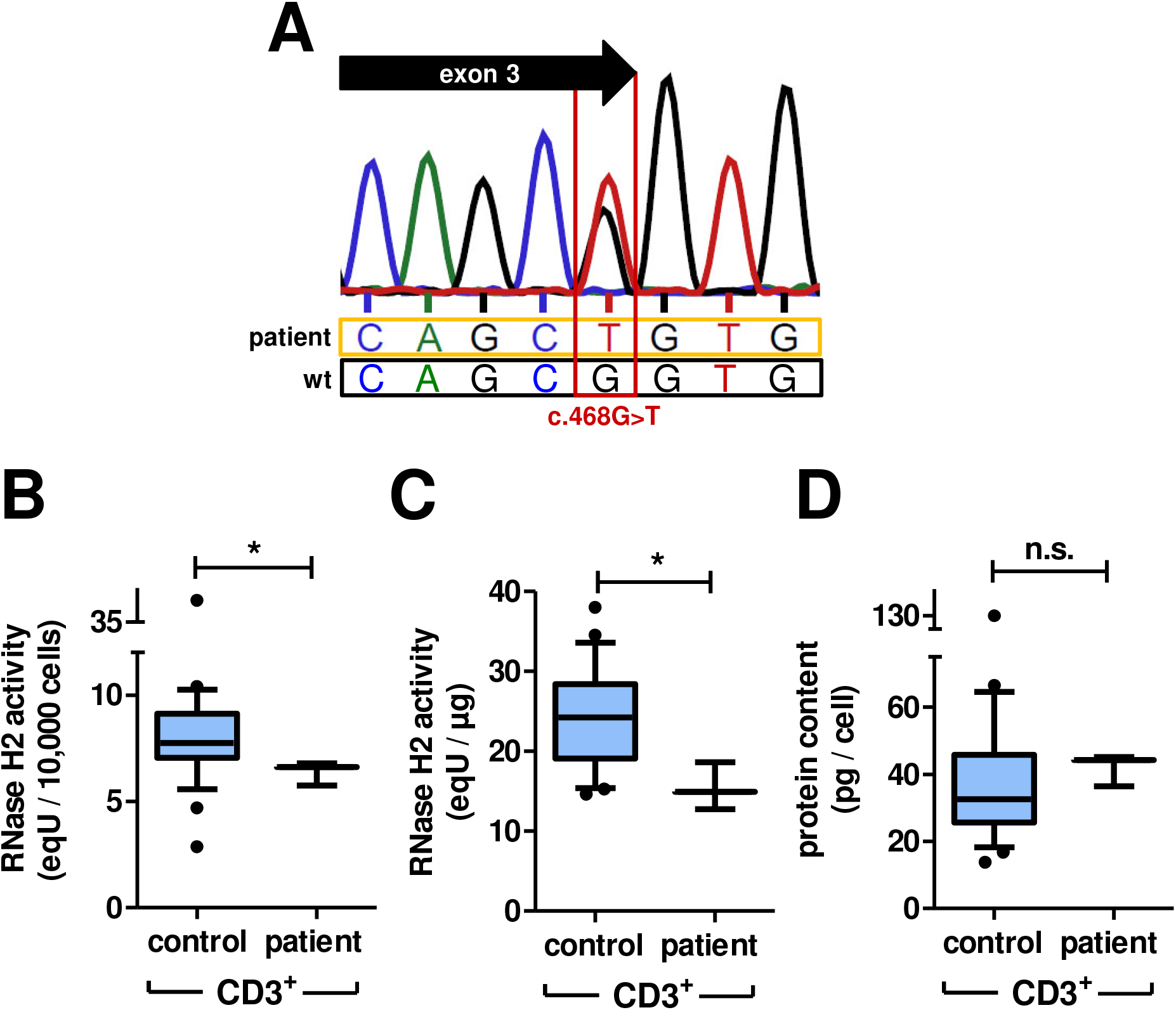
RNase H2 activity in T cells from a patient carrying a heterozygous *RNASEH2C c*.468G>T variant. **A)** Verification of the mutation by Sanger sequencing of DNA from PBMCs. The mutation is silent but affects the last base of exon 3, which is part of the splice donor site. This leads to aberrant splicing of the RNASEH2C mRNA, resulting in a sterile transcript ^22^. **B – D)** RNase H2 activity normalized to cell number (B) or to total cellular protein (C) and total cellular protein content (D) of T cells heterozygous for the *RNASEH2C c*.*468G>T* variant compared to T cells from healthy controls (with unknown *RNASEH2* genotype, n=24). Median is shown, boxes indicate 25^th^ and 75^th^ percentile, whiskers indicate 10^th^ and 90^th^ percentile, extreme outliers (>95^th^ percentile oder < 5^th^ percentile) are shown in the diagram, but were excluded for significance testing via unpaired two tailed t-test with Welch’s correction, **** p < 0.0001, *** p < 0.001, ** p < 0.01 * p < 0.5, n.s. not significant.

## Discussion

We present standardization and validation of an assay allowing for quantification of RNase H2 activity in cell lysates. The assay is based on the experimental procedure published by Crow et al ^42^, and is suitable for screening of clinical samples. Measurement of RNase H2 activity has primarily been performed on recombinant wild type or mutant RNase H2 ^6,12,35,36,54^. Assays based on recombinant protein have important limitations as effects of e.g. protein stability or protein-protein interaction are not captured. Recombinant protein expression requires sequencing and cloning of gene variants, precluding adaptation to settings of clinical screening. Assays involving overexpression of recombinant proteins also do not address effects of intracellular expression levels of functional enzyme. We directly measure RNase H2 activity of cell lysates, enabling determination of enzyme activity per cell or per amount of cellular protein, capturing any effect on levels of functional enzyme in the cell, including alteration of transcription, posttranscriptional regulation, posttranslational modifications and protein stability.

While the assay is based on lymphocytes sorted by FACS in the present study, less sophisticated methods of cell separation are clearly sufficient. In contrast to flow cytometric sorting, immunomagnetic cell separation ^55^, is cost- and labor-efficient, requires only simple equipment, and allows fast enrichment of B or T cells from peripheral blood samples to high purity. Future establishment of a larger and more heterogeneous control cohort evenly distributed between all age groups and gender is necessary to identify potential confounders (e.g. ethnicity, hormonal changes, medication, stress, circadian rhythm, CD4+/CD8+ ratio, etc.) contributing to *inter-* and *intra*-individual variability. While the assay enables reliable detection of activity reduction by 30% comparing triplicate measurements of a single sample to the control group mean (24 individuals), increasing replicate measurements of patient samples to 5 and control cohort size to 100 would improve the minimal detectable activity reduction to 20%, which is in the range of inter-individual variability. Therefore, an activity reduction of less than 20% is unlikely to be clinically relevant. Measurement of RNase H2 activity in triplicate T cell samples from a patient suffering from systemic sclerosis revealed significantly reduced activity by normalization to cell number or to amount of total cellular protein. Since RNase H2 is a nuclear protein and genomic DNA is its substrate, and cell volume the activity per genome, i.e. per cell, seems to be the most important parameter, while cellular volume and protein content are subject to fluctuations, e.g. depending on the cell cycle (Supplemental Figure S2). The assay is fully standardized and externally validated quality controls meeting all requirements for certified reference materials were implemented ensuring inter-laboratory reproducibility. High sensitivity, robustness against impact of sample storage or freezing and a broad working range enable versatile applications.

Collectively, we provide a fully standardized, validated and benchmarked assay suitable for quantification of RNase H2 enzyme activity in clinical cell samples. The assay is sensitive and precise. It revealed differences in RNase H2 activity dependent on cell type and cell cycle phase as well as the reduction of enzyme activity caused by a heterozygous RNASEH2C partial loss-of-function mutation. The assay will be valuable for screening clinical entities for alterations of RNase H2 function in autoimmunity and cancer.

## Acknowledgements

We thank Barbara Utess, Livia Schulze, Christina Hiller and Christa Haase for excellent technical support, Björn Hiller for help with the isolation of primary cells. iPS cells were kindly provided by Michael Haase, Department of Pediatric Surgery, Medical Faculty, TU Dresden. This work was funded by the Deutsche Forschungsgemeinschaft (DFG, German Research Foundation) – Project-ID 369799452 – TRR237 Nucleic Acid Immunity, project B17 to A.R., project B19 to R.B. and project B20 to C.G., as well as by Else Kröner-Fresenius-Foundation grant 060_380627 to M.S.

## Supplemental tables

**Table S1.** RNaseH2 assay working range for different cell types. Columns show minimum, maximum and recommended amount of total protein for each cell type to achieve minimum (8.1 eqU), maximum (197 eqU) or optimal (> 20 eqU, < 100 eqU) substrate conversion rates. The approximate number of cells needed for a yield of 1 μg of total protein is shown on the right.

|  | cell type | minimum<br>amount<br>of<br>protein/<br>well | maximum<br>amount<br>of<br>protein/<br>well | optimal<br>amount of<br>protein/<br>well | number of<br>cells<br>yielding 1<br>µg total<br>protein |
| --- | --- | --- | --- | --- | --- |
| **murine** | CD4<sup>+</sup> primary<br>T cells | 1.1 | 26.2 | 2.7 – 13.3 | 2.6E+04 |
|  | CD11b<sup>+</sup><br>spleen<br>macrophages | 2.9 | 70.6 | 7.2 – 35.8 | 8.2E+03 |
|  | CD11b<sup>+</sup>/CD1<br>1c<sup>+</sup> spleen<br>dendritic cells | 2.8 | 67.2 | 6.8 – 34.1 | 5.8E+03 |
|  | CD19<sup>+</sup><br>primary B<br>cells | 1.3 | 32.6 | 3.3 – 16.6 | 9.7E+04 |
|  | F4/80<sup>+</sup><br>peritoneal<br>macrophages | 1.7 | 42.1 | 4.3 – 21.3 | 8.2E+03 |
|  | CD49f<sup>+</sup><br>epidermal<br>stem cells | 2.3 | 55.3 | 5.6 – 28.1 | 2.0E+04 |
|  | murine<br>embryonic<br>fibroblasts<br>(MEFs) | 8.6 | 210.2 | 21.3 –<br>106.7 | 1.4E+04 |
| **human** | BJ cell line | 32.1 | 782.4 | 79.4 –<br>397.1 | 2.4E+03 |
|  | HeLa cell line | 0.9 | 23.1 | 2.3 – 11.7 | 4.3E+03 |
|  | HEK293T<br>cell line | 0.6 | 15.8 | 1.6 – 8.0 | 4.4E+03 |
|  | CD3<sup>+</sup> primary<br>T cells | 0.5 | 13.4 | 1.4 – 6.8 | 5.4E+04 |
|  | CD19<sup>+</sup><br>primary B<br>cells | 1.5 | 37.1 | 3.8 – 18.8 | 9.4E+03 |
|  | iPS cell line | 0.5 | 13.1 | 1.3 – 6.7 | 1.0E+05 |

## Descriptions for supplemental figures

**Figure S1.**
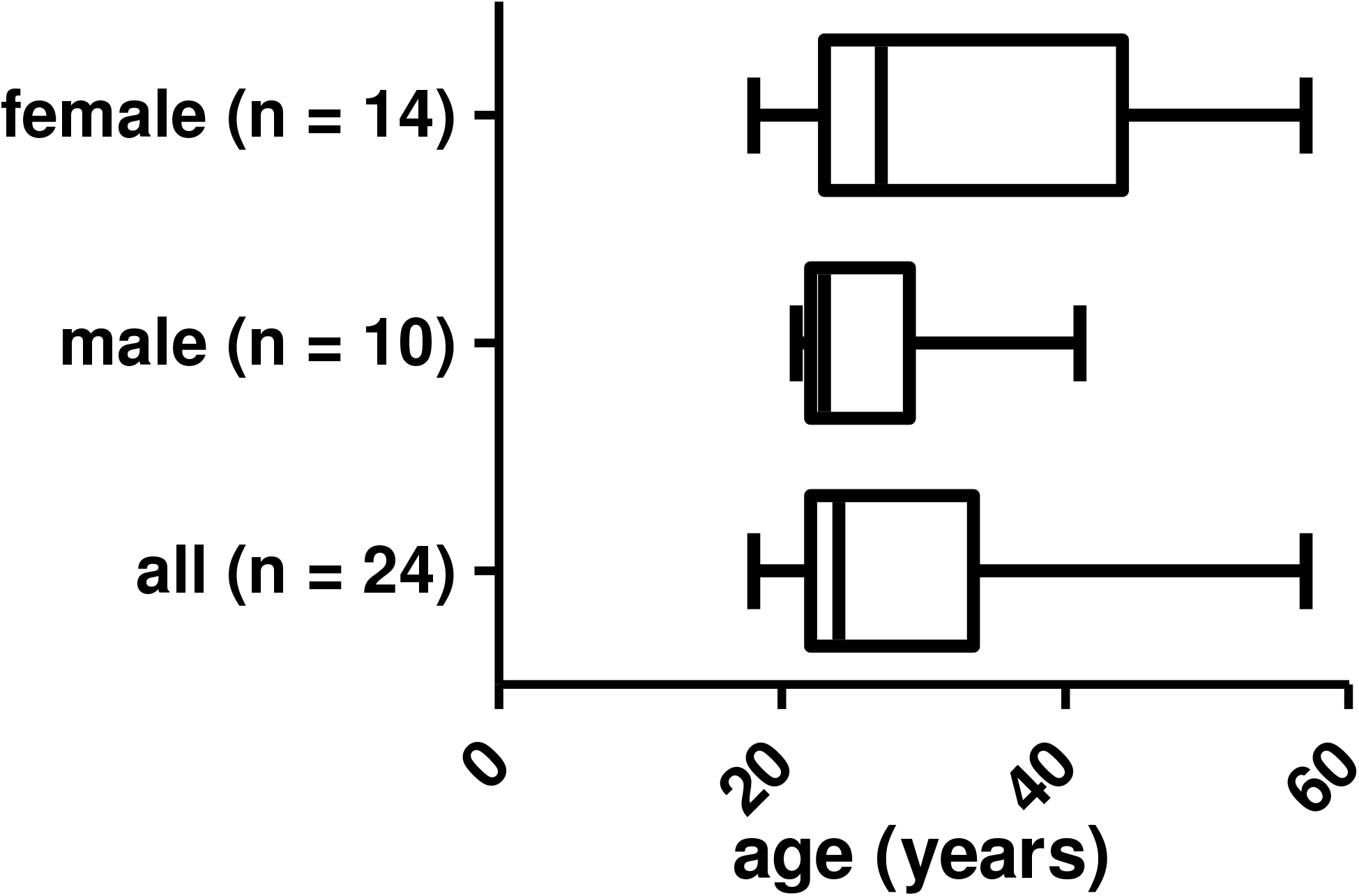
Age and gender of the healthy control group. (24 individuals with unknown RNASEH2 genotype). Median is shown, boxes indicate 25^th^ and 75^th^ percentile, whiskers indicate minimum and maximum.

**Figure S2.**
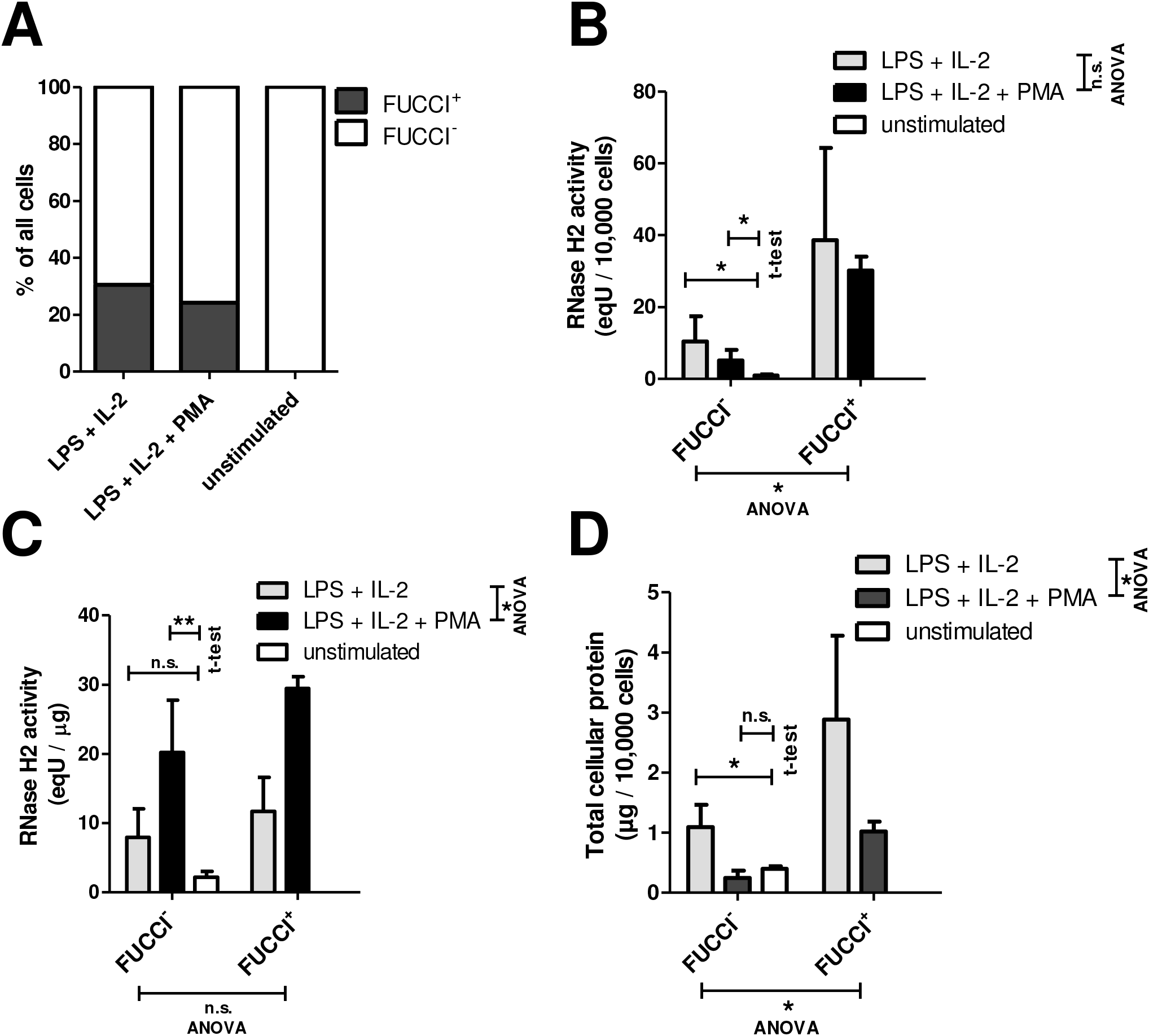
Influence of cell cycle phase on RNase H2 activity. FUCCI cell cycle reporter transgenic mice report cells in S / G2 / M phase of the cell cycle ^58^. Total spleen cells of three FUCCI mice were isolated and stimulated in complete B cell medium for 48 hours with LPS (25 μg / ml) + IL-2 (180 U / ml), LPS (12.5 μg / ml) + IL-2 (180 U / ml) + PMA (5 ng / ml) or left untreated. Between 1×10^5^ and 2×10^6^ FUCCI^+^ and FUCCI-negative CD19^+^ B cells per mouse were sorted by FACS using anti-murine CD19 (eBio1D3) PE antibodies and RNase H2 activity was measured in triplicates. **A)** Fraction of FUCCI^+^ cells upon stimulation. **B)** RNase H2 activity normalized to cell number in B cells in S / G2 / M-phase (FUCCI^+^) versus FUCCI-negative B cells. **C)** RNase H2 activity normalized to total cellular protein in B cells in S /G2 /M-phase (FUCCI^+^) versus FUCCI-negative B cells. **D)** Amount of protein per cell in B cells in S /G2 /M-phase (FUCCI^+^) versus FUCCI^-^ B cells. Cells stimulated with LPS and IL-2 possessed significantly more protein per cell than cells stimulated with LPS, IL-2 and PMA. FUCCI+ cells possessed significantly more cellular protein per cell than FUCCI-negative cells. Increase of cellular protein from FUCCI-negative to FUCCI+ cells was about 2.6-fold. Only stimulation with LPS + IL-2 significantly increased the amount of total cellular protein in FUCCI-negative cells. Mean ± SD is shown, significance was tested via unpaired two tailed t-test, or 2-way ANOVA, as indicated in the graphs, **** p < 0.0001, *** p < 0.001, ** p < 0.01 * p < 0.5, n.s. not significant.

## Notes

*Financial Disclosure and Conflicts of Interest:* The authors declare that there are no conflicts of interests.

### Competing Interest Statement

The authors have declared no competing interest.

